# NADPH oxidase generated reactive oxygen species and aquaporin conduits mediate activity-regulated dendritic plasticity

**DOI:** 10.1101/2020.11.16.384487

**Authors:** Serene Dhawan, Philip Myers, Matthias Landgraf

## Abstract

Neurons utilize plasticity of dendritic arbors as part of a larger suite of adaptive plasticity mechanisms. This explicitly manifests with motoneurons in the Drosophila embryo and larva, where dendritic arbors are exclusively postsynaptic and are used as homeostatic devices, compensating for changes in synaptic input through adapting their growth and connectivity. We recently identified reactive oxygen species (ROS) as novel plasticity signals instrumental in this form of dendritic adjustment. ROS correlate with levels of neuronal activity and negatively regulate dendritic arbor size. Here, we investigated NADPH oxidases as potential sources of such activity-regulated ROS and implicate Dual Oxidase (but not Nox), which generates hydrogen peroxide extracellularly. We further show that the aquaporins Bib and Drip, but not Prip, are required for activity-regulated ROS-mediated adjustments of dendritic arbor size in motoneurons. These results suggest a model whereby neuronal activity leads to activation of the NADPH oxidase Dual Oxidase, which generates hydrogen peroxide at the extracellular face; aquaporins might then act as conduits that are necessary for these extracellular ROS to be channeled back into the cell where they negatively regulate dendritic arbor size.

## 1 Introduction

Neurons are inherently plastic and their ability to respond to changes in synaptic transmission or activity patterns is central to many processes, from learning and memory (Martin et al., 2000; Stuchlik, 2014) to homeostatic adjustments that stabilize circuit function (Turrigiano and Nelson, 2000; Pozo and Goda, 2010; Wefelmeyer et al., 2016; Frank et al., 2020). We recently identified reactive oxygen species (ROS) as novel signals required for activity-regulated plasticity. ROS have long been known to affect neuronal development and function. Commonly associated with pathological conditions, ageing and disease, the roles of ROS as signalling molecules under normal physiological conditions are much less well understood (Milton et al., 2011; Oswald et al., 2018b; Peng et al., 2019). At present, we know that hydrogen peroxide (H_2_O_2_) is the species predominantly required for homeostatic maintenance of synaptic transmission at the neuromuscular junction in *Drosophila* larvae, and for adaptive structural changes of synaptic terminal arbors following periods of over-activation. H_2_O_2_ is also sufficient to induce structural changes that largely phenocopy effects of over-activation, suggesting that ROS mediate plastic adjustments downstream of neuronal activity (Oswald et al., 2018a). Given that neuronal activity has a high energetic cost, correlating with metabolic demand (Attwell and Laughlin, 2001; Zhu et al., 2012), a simplistic working model has been proposed whereby ROS, largely generated as obligate by-products of aerobic metabolism (Halliwell, 1992), regulate plasticity by providing ongoing feedback on a neuron’s activation status (Hongpaisan et al., 2003, 2004; Oswald et al., 2018a).

In addition to mitochondria NADPH oxidases are also well documented sources of activity-generated ROS (Hongpaisan et al., 2004; Massaad and Klann, 2011; Baxter and Hardingham, 2016; Hidalgo and Arias-Cavieres, 2016). Here, we focus on NADPH oxidases, a family of differentially expressed multisubunit enzymes (Nox 1-5 and Dual Oxidase 1-2) that catalyse the transfer of an electron from cytosolic NADPH to oxygen to generate ROS at the extracellular face of the plasma membrane (Lambeth, 2002; Panday et al., 2015). NADPH oxidases are commonly associated with immune responses, but may also regulate aspects of nervous system development such as neuronal polarity (Wilson et al., 2015), growth cone dynamics (Munnamalai and Suter, 2009; Munnamalai et al., 2014), and intriguingly, synaptic plasticity (Tejada-Simon et al., 2005).

Using the *Drosophila* locomotor network as an experimental model, we focused on the regulation of dendritic growth of identified motoneurons, a sensitive assay of structural plasticity, whereby neuronal activity and associated ROS reduce dendritic arbor size (Oswald et al., 2018a). Based on single cell-specific targeting of RNAi knockdown constructs, our data suggest that the NADPH oxidase Dual Oxidase (Duox), but not Nox, is required for activity-regulated generation of ROS. Because Duox generates H_2_O_2_ at the outer face of the plasma membrane, we tested a requirement for aquaporin channels as conduits transporting extracellular ROS into the cytoplasm, as had been shown in other systems (Miller et al., 2010; Bertolotti et al., 2013). Indeed, we found a requirement for two of three characterised *Drosophila* aquaporins, Bib and Drip (but not Prip). Overall, our data suggest that neuronal activity promotes Duox-mediated generation of extracellular H_2_O_2_, returned into the cytoplasm via aquaporin channels, where it inhibits dendritic growth. Moreover, our findings imply that Duox-generated H_2_O_2_ additionally acts non-autonomously on neighbouring synaptic terminals.

## 2 Methods

### 2.1 Fly genetics

*Drosophila melanogaster* strains were maintained on a standard apple juice-based agar medium at 25°C. The following fly strains were used: *OregonR* (#2376, Bloomington Drosophila Stock Center), *UAS-dTrpA1* in attP16 *(Hamada et al., 2008), UAS-Duox.RNAi* (#32903, BDSC), *UAS-Nox.RNAi* (Ha et al., 2005; FBal0191562), *UAS-bib.RNAi (I)* (#57493, BDSC), *UAS-bib.RNAi (II)* (#27691 BDSC), *UAS-Drip.RNAi (I)* (#44661, BDSC), *UAS-Drip.RNAi (II)* (#106911, Vienna *Drosophila* Resource Centre), *UAS-Prip.RNAi (I)* (#50695, BDSC), *UAS-Prip.RNAi (II)* (#44464, BDSC), and *UAS-secreted human-catalase* (Ha et al., 2005). Transgene expression was targeted to 1-3 RP2 motoneurons per nerve cord using a stochastic FLPout strategy, as detailed previously (Fujioka et al., 2003; Ou et al., 2008). Briefly, yeast Flippase expression is directed to RP2 motoneurons at embryonic stages only (at a lower level also to the aCC motoneuron and pCC interneuron) via RN2-FLP ((Fujioka et al., 2003; Ou et al., 2008)), then initiates permanent GAL4 expression in a subset of RP2 motoneurons via the FLP-conditional driver, *tub84B-FRT-CD2-FRT-GAL4* (Pignoni and Zipursky, 1997).

### 2.2 Larval staging and dissection

Eggs were collected at 25°C over an 8-hour period on an apple juice-based agar medium supplemented with a thin film of yeast paste and, following continued incubation at 25°C, were subsequently screened for freshly hatched larvae, selected against the presence of fluorescently marked (*deformed-GMR-YFP*) balancer chromosomes. Larvae were transferred to a fresh agar plate with yeast paste, incubated at 25°C (aquaporin/catalase experiments) or 27°C (NADPH oxidase experiments) and allowed to develop to the third instar stage (approximately 48 hours after larval hatching), followed by dissection in external saline (pH 7.15) (Marley and Baines, 2011). Nerve cords were transferred with a BSA coated glass capillary onto a poly-L-lysine coated (Sigma-Aldrich) cover glass (22×22mm), positioned dorsal side up. A clean cover glass was placed on top with two strips of electrical tape used as spacers.

### 2.3 Image acquisition

Nerve cords were imaged within a 5 minute window from dissection using a custom-built spinning disk confocal microscope consisting of a CSU-22 field scanner (Yokagawa), mounted on a fixed stage upright Olympus microscope frame (BX51-WI), equipped with a single objective piezo focusing device (Physik Instruments), a 60x/1.2 NA water immersion objective (Olympus), external filter wheel (Sutter) and programmable XY stage (Prior). Images were acquired at an effective voxel size of 0.217 × 0.217 × 0.3 μm using a back-thinned Evolve EMCCD camera (Photometrics), operated via MetaMorph software (Molecular Devices).

### 2.4 Neuron Reconstruction

Dendritic arbor reconstructions were carried out in Amira 6.5 (FEI). A deconvolution algorithm was used to reassign photons from out-of-focus optical sections to their points of origin, thus improving the signal-to-noise ratio of the image stack. Subsequently, thresholding of voxel grey values was used to segment the fluorescent arbor from background. Structures, which did not require reconstruction, i.e. the cell body and primary neurite, were manually removed. Post segmentation, the Amira automatic reconstruction algorithm was used to convert the centrelines of the user-defined segmentation into a spatial graph structure. This structure was manually reviewed and edited to correct for ‘loops’ and other artefacts of the automatic reconstruction process. Quantification of cell body area was conducted in ImageJ (National Institutes of Health) by manually tracing around individual cell bodies.

### 2.5 Data handling and statistical analysis

All data handling and statistical analyses were carried out in R. A Shapiro-Wilk test was used to confirm normality of all dendritic arbor reconstruction data presented. A one-way analysis of variance (ANOVA) followed by Tukey’s multiple comparisons test was used to compare experimental manipulations to the controls where *p<0.05, **p<0.01, ***p<0.001 and ****p<0.0001.

## 3 Results

### 3.1 ROS generated by Dual oxidase are required for activity-dependent adjustments of dendritic arbor size and geometry

We have previously shown that the postsynaptic dendritic arbor behaves as a homeostatic device, adjusting its size and geometry in an activity-dependent manner to facilitate receipt of an appropriate level of presynaptic input (Tripodi et al., 2008; Oswald et al., 2018a). Moreover, we have identified ROS signalling as necessary and sufficient for this structural remodelling to occur (Oswald et al., 2018a). Building on this work, here we set out to investigate whether NADPH oxidases might play a role during activity and ROS-regulated structural plasticity. Conveniently, *Drosophila* codes for only two NADPH oxidases; *dDuox*, an orthologue of vertebrate dual oxidase, and *dNox*, which is closely related to human Nox5 (Kawahara et al., 2007). To investigate if either or both contributed to activity-regulated structural plasticity of dendrites, we targeted the expression of RNAi constructs designed to knockdown *dDuox* or *dNox* to the well-characterized ‘RP2’ motoneuron (Sink and Whitington, 1991; Baines et al., 1999; Landgraf et al., 2003), with and without concomitant overactivation. We then analysed RP2 dendritic arbors morphometrically to quantify the extent to which these manipulations impacted their development.

In accordance with previous findings (Oswald et al., 2018a), neuronal overactivation by targeted dTrpA1 misexpression in individual RP2 motoneurons resulted in significantly smaller dendritic arbors with reduced dendritic length and branch point number, as compared to non-manipulated controls (Figure 1A-C). However, co-expression of an RNAi construct designed to knockdown *dDuox* along with *UAS-dTrpA1* significantly attenuated the activity-induced reduction of dendritic length and branch point number. In contrast, the RNAi-mediated knockdown of *dNox* had no discernible effect on arbor morphology. Mis-expression of *UAS-dDuox.RNAi* or *UAS-dNox.RNAi* transgenes alone under endogenous activity conditions, i.e. in the absence of dTrpA1 manipulation, produced no significant differences in arbor characteristics between all genotypes (Figure 1A-C).

**Figure 1.**
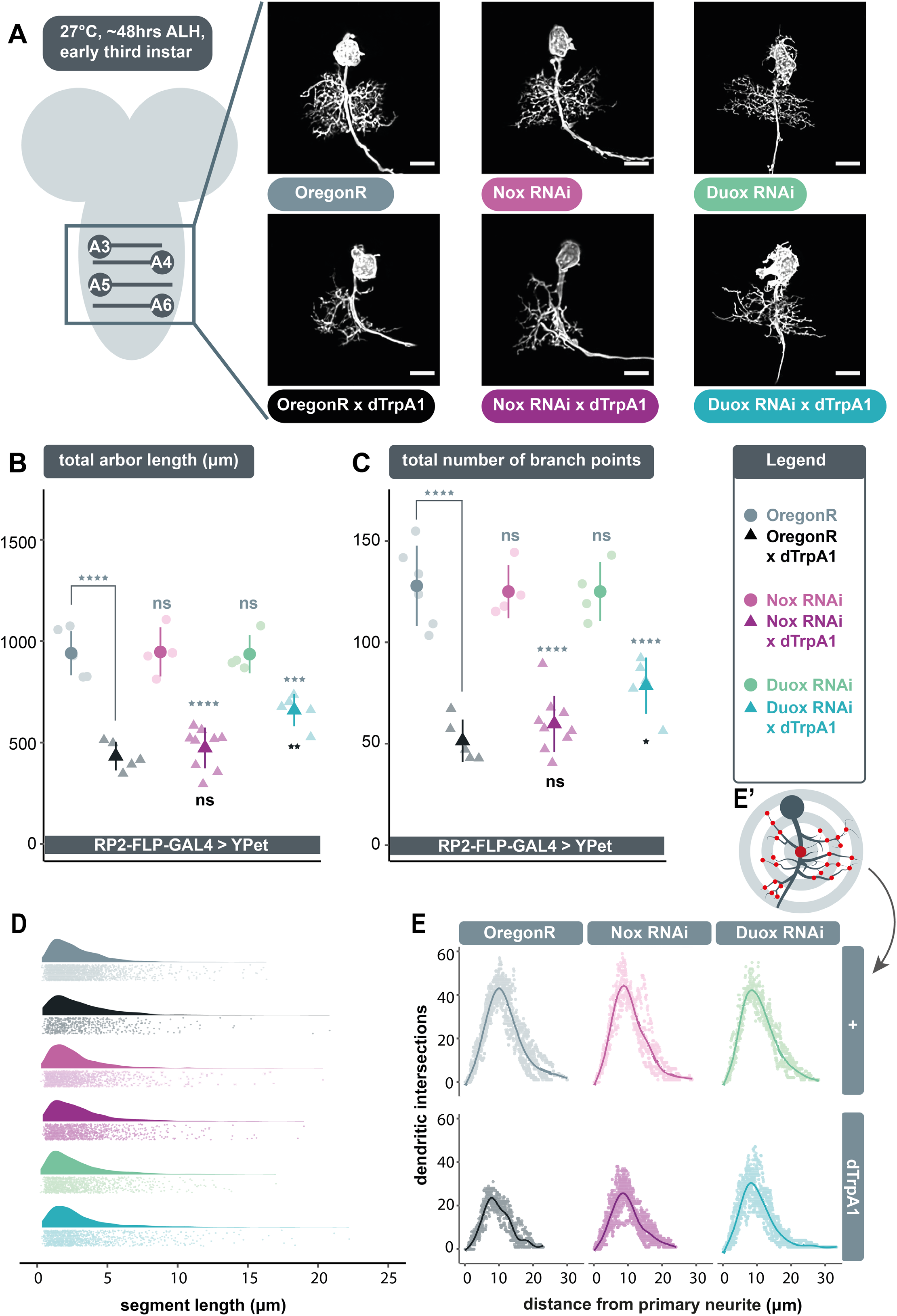
Extracellular ROS generated by *dDuox*, but not *dNox*, are required for homeostatic structural plasticity in response to increased neuronal activity. **(A)** Maximum intensity z-projections of representative RP2 motoneurons located within abdominal segments A3-6 in the ventral nerve cord, from young third instar larvae raised at 27°C and dissected 48 hours after larval hatching. Scale bars: 15μm. **(B-C)** Targeted expression of *UAS-dDuox-RNAi*, but not *UAS-dNox-RNAi*, significantly suppresses activity-induced reduction in total dendritic arbor length and number of branch points relative to *UAS-dTrpA1* manipulated motoneurons. In absence of *UAS-dTrpA1,* RP2 dendrites expressing these RNAi constructs are comparable to controls (ANOVA, ns = not significant, *p<0.05, **p<0.01, ***p<0.001, ****p<0.0001). Comparisons with non-manipulated controls are shown directly above data points (light grey) and comparisons with the *UAS-dTrpA1* overactivation condition are shown directly below (black). **(D)** Irrespective of neuronal activity regime, expression of *UAS-dDuox-RNAi* or *UAS-dNox-RNAi* does not alter the frequency distribution of dendritic segment lengths. A segment is defined as the distance between two branch points, or, in the case of a terminal neurite, a branch point and the tip. Frequency density plot shown in darker color, with all individual data points plotted below in a corresponding lighter shade. **(E)** Sholl analyses indicate that genetic manipulations do not obviously change the relative distribution of dendrites in 3D space. Mean dendritic intersections as a function of distance from the midpoint of the primary neurite **(E’)** are shown as a solid coloured line, all individual data points are shown in a corresponding lighter shade.

Next, we asked what changes in arbor structure might lead to these activity-regulated differences. We considered two principal possibilities, namely changes in the pattern of growth versus changes in the number of dendritic segments generated. Our analysis of the underlying frequency distribution of dendritic segment lengths points to the latter. Across genotypes, segment lengths were unimodally distributed with a peak at ~2μm that sharply tailed off and became negligible around 8-9μm (Figure 1D). This indicates that the reduction of RP2 motoneuron dendritic arbors caused by dTrpA1-mediated overactivation is the result of fewer dendritic segments formed or maintained, as opposed to changes in the fundamental mode of arbor elaboration, which may lead to the generation of similar numbers of shorter dendritic segments. Secondly, this analysis suggests that the partial attenuation of the overactivation phenotype, conferred by cell-specific *UAS-Duox.RNAi* expression, is produced by increasing dendritic segment number.

We also examined dendritic topography via Sholl analysis to see if changes in arbor length might impact on the ability to invade different regions of the neuropil. The origin of the concentric Sholl spheres was defined as the midpoint on the primary neurite (from which all dendrites arise) between the first and last branch points (Figure 1E’). Irrespective of genotype, the majority of arbor length was concentrated just under halfway along the arbor’s expanse with respect to the primary neurite (Figure 1E). This suggests that the genetic manipulations conducted do not obviously lead RP2 motoneurons to alter the placement or density of their dendritic segments. This is in agreement with earlier findings of motoneurons being specified by genetically separable programmes for dendritic growth and dendritic positioning (Ou et al., 2008)

In summary, these data implicate dDuox, but not dNox, as necessary for implementing structural plasticity of dendritic arbors in response to elevated activity.

### 3.2 Aquaporin channel proteins Bib and Drip mediate activity-and ROS-dependent neuronal plasticity

NADPH oxidases generate ROS extracellularly, which poses the question of how such NADPH-generated ROS affect dendritic growth. Do they do so by modifying extracellular components or via intracellular events, which would require entry into the cell. We focused on investigating the latter. Owing to the large dipole moments of key ROS like H_2_O_2_, simple diffusion across the hydrophobic plasma membrane, as seen with small and non-polar molecules, is limited. Instead, evidence from other experimental systems points towards a model of facilitated diffusion involving aquaporins (Bienert et al., 2007; Miller et al., 2010; Bertolotti et al., 2013). Classically, the importance of these channel proteins has been stressed in the process of transmembrane fluid transport. However, some lines of research suggest that aquaporins can regulate the downstream signalling pathways that rely on ROS as a second messenger, by controlling entry of ROS into the cytosol (Miller et al., 2010; Bertolotti et al., 2013).

We postulated that following neuronal overactivation extracellular ROS generated by Duox are brought into the cell via aquaporin channels, where they can then trigger compensatory structural changes in dendritic arbor size. To test this model, we overactivated individual RP2 motoneurons by targeted expression of dTrpA1 whilst simultaneously expressing independently generated RNAi constructs designed to knockdown genes that encode aquaporin channels: *big brain* (*bib*), *Drosophila* i*ntrinsic proteins* (*Drip*) or *Pyrocoelia rufa integral proteins* (*Prip*). Under conditions of cell-autonomous neuronal overactivation, targeted co-expression of *UAS-bib.RNAi* or *UAS-Drip.RNAi* transgenes resulted in a significant ‘rescue’ of dendritic arbor length and branch point number, i.e. abrogation of activity-induced arbor reduction, similar to the effects of *UAS-dDuox.RNAi* (Figure 2B-C) as compared to dTrpA1 manipulated RP2 motoneurons of similar age (48hrs ALH at 25°C). In contrast, targeted expression of *UAS-Prip.RNAi* transgenes did not have any significant effect on arbor morphology.

**Figure 2.**
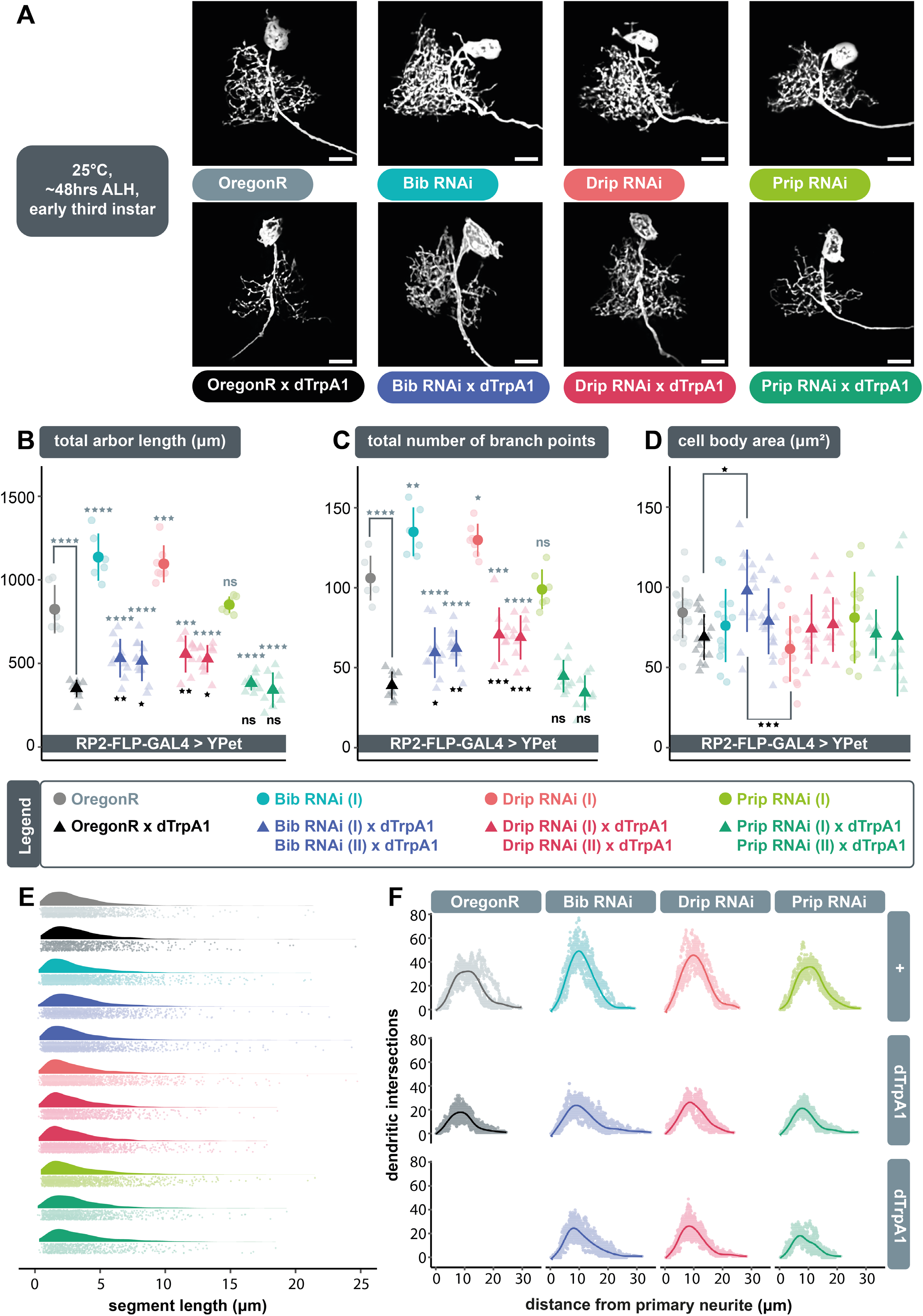
The aquaporins encoded by *bib* and *Drip*, but not *Prip*, are necessary for ROS and activity-induced dendritic plasticity. **(A)** Maximum intensity z-projections of representative RP2 motoneurons from young third instar larvae raised at 25°C and dissected 48 hours after larval hatching. Scale bars: 15μm. **(B-C)** Targeted expression of *UAS-bib-RNAi* or *UAS-Drip-RNAi*, but not *UAS-Prip-RNAi* in RP2 motoneurons significantly attenuates the reduction in dendritic arbor length and number of branch points caused by dTrpA1-mediated neuronal overaction. Two independently generated RNAi constructs were tested for each aquaporin. Expression of *UAS-bib-RNAi* or *UAS-Drip-RNAi* without concomitant dTrpA1 manipulation produces a dendritic overgrowth phenotype characterized by increased arbor length and branching complexity relative to non-manipulated controls. **(D)** Quantification of cell body size, measured as the maximum area of the 2D cross-section through the centre of the soma, shows no consistent differences between controls and *UAS-aquaporin RNAi* manipulations. **(B-D)** ANOVA, ns = not significant, *p<0.05, **p<0.01, ***p<0.001, ****p<0.0001. Comparisons with non-manipulated controls are shown directly above data points (light grey) and comparisons with overactivated controls are directly below (black). **(E)** Expression of RNAi transgenes designed to knock down specific aquaporins does not alter fundamental arbor structure under conditions of endogenous activity or chronic overactivation. Frequency density plot of dendritic segment lengths shown in darker color, with all individual data points plotted below in a corresponding lighter shade. **(F)** The branching topology of RP2 motoneurons is not affected by *UAS-aquaporin-RNAi* manipulations. Mean dendritic intersections as a function of distance from the midpoint of the primary neurite are shown as a solid coloured line, all individual data points are shown in a corresponding lighter shade.

Intriguingly, mis-expression of *UAS-bib.RNAi* or *UAS-Drip.RNAi* alone, without concomitant dTrpA1 activity manipulation, produced a dendritic overgrowth phenotype, while targeted *UAS-Prip*-*RNAi* expression had no measurable impact on dendritic development. Relative to non-manipulated control neurons, neurons expressing *UAS-bib.RNAi* or *UAS-Drip.RNAi* exhibited a significant increase in dendritic length (Figure 2B) and branching complexity (Figure 2C) in the ventral part of their arbor, comparable to what one normally observes in wild-type RP2 motoneurons at later larval stages. As seen with *UAS-dDuox.RNAi* manipulation, here too changes in dendritic growth appear to result from changes in the number rather than length of dendritic segments generated (Figure 2E). A Sholl analysis also did not report significant differences in the relative distribution of dendritic segments within the neuropil, indicating that RP2 motoneurons target their normal neuropil territories irrespective of aquaporin knockdown manipulation (Figure 2F).

We wondered whether such overgrowth might be caused by increased internal osmotic pressure, as a result of impaired aquaporin function, which in turn may have stimulated mechanically sensitive proteins expressed within the neuronal growth cone (Kerstein et al., 2015). We reasoned that increased osmotic pressure would result in dilation of the soma. However, we could not see a consistent pattern maximum cell body area being affected by expression of *UAS-aquaporin.RNAi* transgenes (Figure 2D).

In summary, these data demonstrate a requirement for the aquaporin channels Bib and Drip, but not Prip, in mediating activity-regulated structural adjustment of dendritic arbor size and geometry. However, since mis-expression of *UAS-bib.RNAi* or *UAS-drip.RNAi* alone (in the absence of dTrpA1-mediated overactivation) leads to a dendritic overgrowth phenotype, it is unclear if aquaporins and neuronal activity act in the same or in separate, parallel pathways that regulate dendritic growth in opposite ways.

### 3.3 Extracellular ROS act as negative regulators of dendritic growth, also under conditions of endogenous activity

The above data suggest that extracellular ROS provide a negative feedback signal, which following dTrpA1-mediated overactivation, facilitates reduction of dendritic arbor size. We wondered if at basal levels of neuronal activity extracellular ROS might also act as negative regulators of dendritic growth. If that was the case, then it could explain why expression of RNAi constructs for knocking down aquaporin channels would have the effect of increased dendritic growth due to decreased influx of extracellular ROS. We reasoned that if correct, then expression of a secreted, extracellular form of catalase, a potent quencher of H_2_O_2_, should lead to increased dendritic growth, similar in effect as expression of *UAS-aquaporin.RNAi* transgenes.

Indeed, we found that expression of *UAS-human-secreted-catalase* in RP2 motoneurons produced a dendritic overgrowth phenotype comparable to that produced by targeted expression of *UAS-bib.RNAi* or *UAS-Drip.RNAi* (Figure 2A,3A). This overgrowth was characterized by dense dendritic arbors with significantly larger total arbor length as compared to non-manipulated controls (Figure 3B). As with other manipulations above, here too the relative distribution of dendritic segments and arbor topography remained comparable to controls (Figure 3E, F). Somewhat unexpected though, our analysis indicates no change in branch point number despite the increase in arbor size (Figure 3C). This is counter-intuitive and we think the most parsimonious explanation for this is artefactual deflation of segment number, caused by the high density of dendritic segments in these overgrown neurons, leading to failure of resolving all during the reconstruction process.

**Figure 3.**
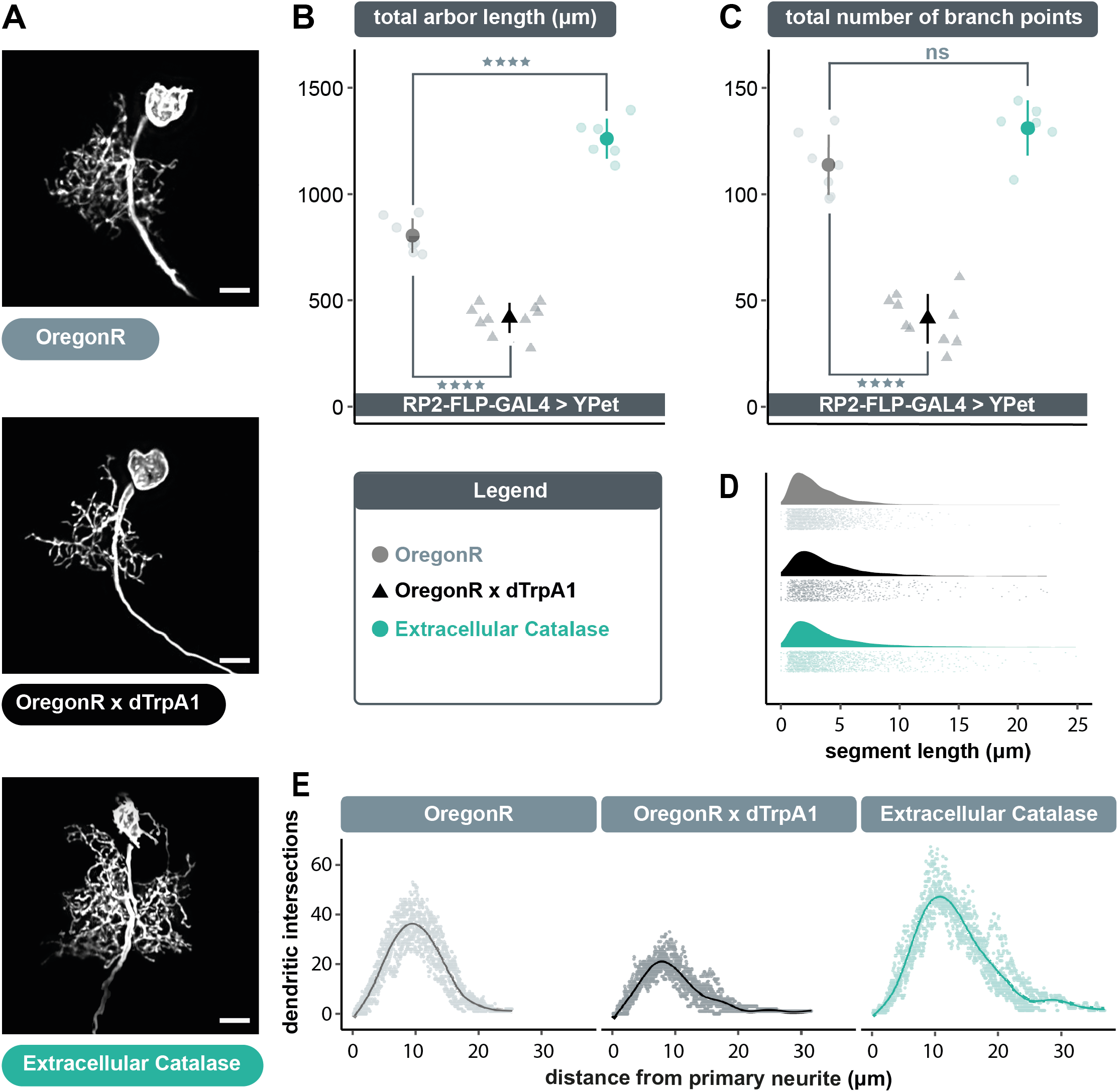
Cell-specific expression of extracellular, human secreted catalase produces a dendritic overgrowth phenotype comparable to that produced by *UAS-bib-RNAi* or *UAS-Drip-RNAi* expression. **(A)** Maximum intensity z-projections of representative RP2 motoneurons, from young third instar larvae raised at 25°C and dissected 48 hours after larval hatching. Scale bars: 15μm. **(B-C)** Targeted expression of *UAS-human secreted catalase* in RP2 motoneurons results in a significant increase or ‘overgrowth’ of dendritic arbor length relative to otherwise non-manipulated neurons, but does not change the total number of branch points (unexpected and likely caused by poor signal-to-noise ratio resulting from very dense branching of these enlarged arbors) (ANOVA, ns = not significant, *p<0.05, **p<0.01, ***p<0.001, ****p<0.0001). **(D)** *UAS-human secreted catalase* expression in RP2 motoneurons does not change the frequency distribution of dendritic segment lengths relative to overactivated or non-manipulated controls Frequency density plot shown in darker color, with all individual data points plotted below in a corresponding lighter shade. **(E)** Sholl analyses indicate that *UAS-human secreted catalase* expression does not obviously cause RP2 motoneurons to alter the placement or density of their dendrites. Mean dendritic intersections as a function of distance from the midpoint of the primary neurite are shown as a solid coloured line, all individual data points are shown in a corresponding lighter shade.

In conclusion, in this study we identified a selective requirement for the NADPH oxidase, Duox (but not Nox), in activity-regulated adjustment of dendritic arbor growth during larval nervous system development. Thus generated extracellular ROS mediate autocrine signalling, that depends on aquaporins Bib and Drip (but not Prip) as conduits for channeling extracellular H_2_O_2_ back into the cell. Overall, extracellular ROS act as negative feedback signals that mediate homeostatic adjustment of dendritic arbor size. That this process also operates at physiological activity levels was revealed by manipulations where we expressed secreted Catalase to quench extracellular ROS in the immediate vicinity of the neuron. Similar to what we find when targeting the aquaporins Bib and Drip for RNAi knockdown, this resulted in enlarged dendritic arbors, as would be predicted when relieving the ROS-brake on dendritic growth.

## 4 Discussion

A prerequisite of flexible yet stable neural circuitry is the ability to detect and appropriately respond to changes in activity, particularly perturbations that push activity towards extremes of quiescence or saturation (Turrigiano and Nelson, 2004; Yin and Yuan, 2014). Here, we focused on structural plasticity of dendrites in response to changes in neuronal activation. Seminal comparative studies in mammals showed that the size and complexity of dendritic arbors correlates with the range and amount of synaptic input received (Purves and Lichtman, 1985; Ivanov and Purves, 1989). Similarly, in the *Drosophila* larval locomotor network we demonstrated that during development the dramatic growth of motoneuron dendritic arbors, which scales with overall body growth, facilitates increases in presynaptic input and thus also of the amount of synaptic drive necessary for appropriate levels of muscle activation (Zwart et al., 2013). This kind of structural plasticity is not particular to periods of growth, but also evident following activity manipulations. For example, changes in the number of active presynaptic sites are compensated for by complementary changes in postsynaptic dendritic arbor size, suggesting that neurons use their dendritic arbors as structural homeostatic devices (Tripodi et al., 2008). Activity-generated ROS are necessary for this plasticity to occur, ROS acting as brakes on dendritic arbor growth or maintenance (Oswald et al., 2018a). Here, we build on this work and identify the NADPH oxidase, Duox, as a key source of such activity-generated ROS. Moreover, the topography of Duox, which is known to reside within the plasma membrane (Morand et al., 2009), is such that it generates H_2_O_2_ into extracellular space (Fogarty et al., 2016), yet our manipulations show that H2O2 is required intracellularly by the cell that has been over-activated (Oswald et al., 2018a). This raised the question of how such extracellular ROS re-enter the neuronal cytoplasm. We provide evidence that the aquaporin channels Bib and Drip, but not Prip, are necessary for activity-regulated dendritic structural remodelling. Specifically, targeting *bib* or *Drip* for RNAi-mediated knockdown in individual RP2 motoneurons partially attenuates the overactivation phenotype of smaller dendritic arbors. These observations suggest that specific aquaporins located in the somatodendritic compartment of motoneurons act as conduits for extracellular H2O2, to enter the cytoplasm and there modulate dendritic growth pathways. Based on our previous observations, we speculate that these extracellular H2O2 act, amongst others, on the cytoplasmic redox-sensitive dimer DJ-1β, which we have shown necessary for structural and physiological changes in response to activity-generated ROS (Oswald et al., 2018a). In addition, the partial penetrance of RNAi knockdown phenotypes, combined with the observation that two distinct aquaporin encoding genes are involved, suggests that there may be functional redundancy between these aquaporins; both are likely expressed and one channel can compensate for the absence of the other owing to similarities in their biochemical activity and substrate specificity. Such redundancy has previously been observed in the tsetse fly, where simultaneous downregulation of multiple aquaporins exacerbates the negative effects on female fecundity produced by individual aquaporin knockdowns (Benoit et al., 2014). Supporting this working model of extracellular ROS acting as negative regulators of dendritic growth, we find that cell-specific expression of RNAi transgenes designed to knockdown *bib* or *Drip* alone produces a dendritic overgrowth phenotype, as would be predicted by reduced influx of extracellular H_2_O_2_ into the cytoplasm. Furthermore, such a model predicts that local quenching of extracellular H_2_O_2_ should have a comparable effect, which indeed we do find.

Altogether, these data are compatible with a model whereby neuronal activity leads to activation of Duox, which generates H2O2 into the extracellular space surrounding the active neuron. Aquaporins then act as conduits that channel these extracellular ROS back into the cytoplasm where they subsequently trigger signalling cascades that negatively regulate dendritic size (Figure 4).

**Figure 4.**
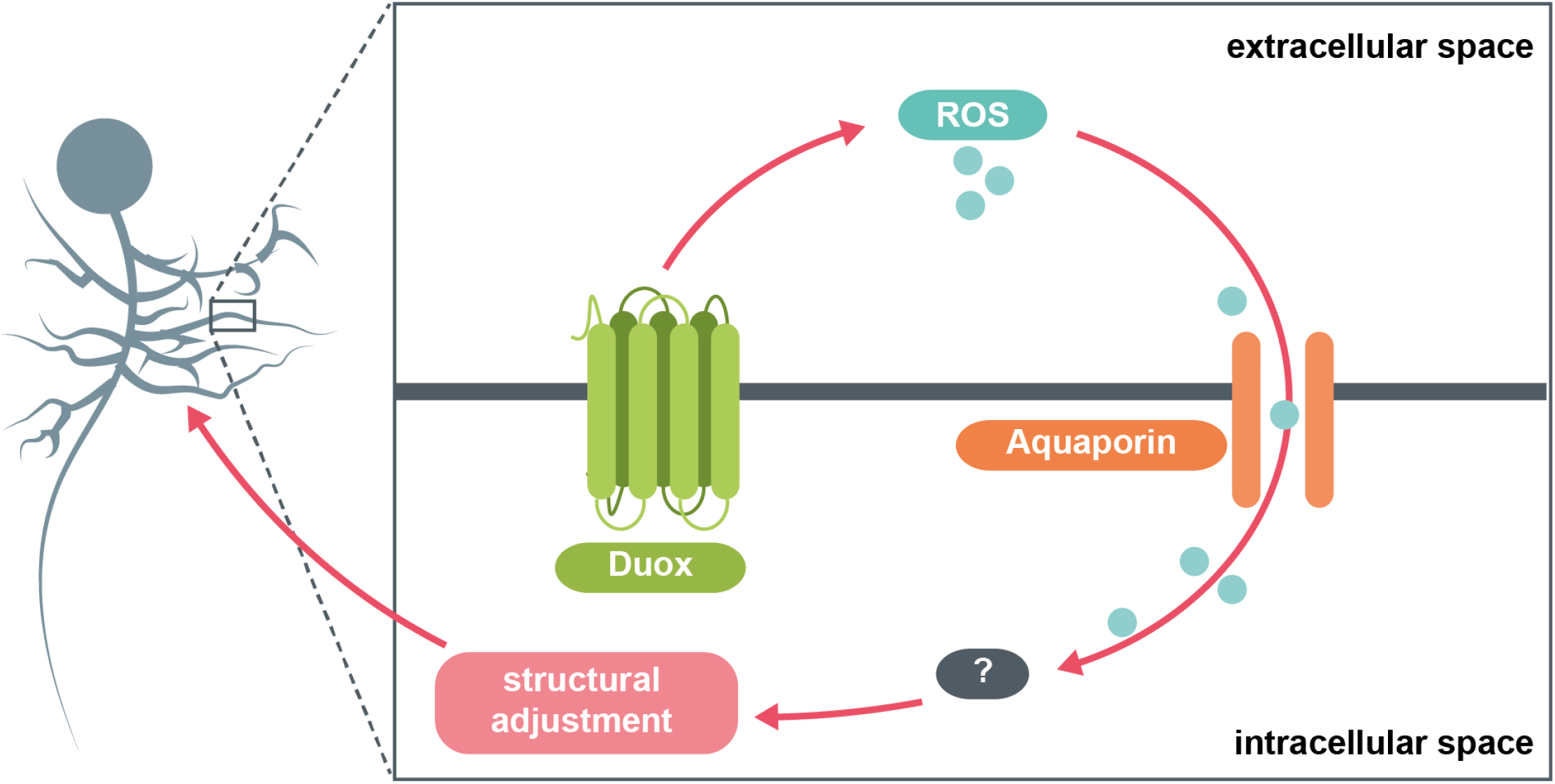
Model summary. The NADPH oxidase Duox generates ROS at the extracellular face of the plasma membrane in response to increases in neuronal activity. These ROS are brought back into the cytosol via specific aquaporin channel proteins. Here, they interact with various intracellular pathways, potentially involving the redox-sensitive dimer DJ-1β, which in turn mediate adaptive reductions in dendritic arbor size.

### 4.1 Extracellular Duox-generated ROS as a potential means for coordinating network-wide homeostatic structural adjustments

Whilst ROS typically operate as intracellular second messengers, (Forman et al., 2014; Schieber and Chandel, 2014), their long-range effects have also been observed during paracrine H_2_O_2_ signalling in vertebrate and invertebrate inflammatory responses (Niethammer et al., 2009; Moreira et al., 2010). In light of this, our finding that extracellular, activity-generated ROS are necessary for structural plasticity raises the intriguing possibility of an intercellular redox-based communication network that coordinates homeostatic structural adjustments more widely, potentially within local volumes. Synaptic clefts in *Drosophila* are approximately 10–20 nm in width (Prokop and Meinertzhagen, 2006). ROS such as O_2_^−^ and H_2_O_2_ have been reported to travel distances in the order of several micrometers within living tissue (Cuypers et al., 2016; Krumova and Cosa, 2016). It follows that extracellular ROS generated by Duox in postsynaptic dendrites could traverse the synaptic cleft and act retrogradely on the surface of presynaptic partner terminals, e.g. on ion channels (Sah et al., 2002; Sesti et al., 2010; Sahoo et al., 2014). Such ROS could further enter the cytosol of presynaptic partners via aquaporin channels and trigger compensatory structural remodelling. In addition, in analogy to retrograde nitric oxide signalling (Hardingham et al., 2013), Duox generated extracellular ROS have the potential to modify pre- and postsynaptic terminals within a local volume of the neuropil, thus acting as regional activity-triggered modulators even between non-synaptic neurons. Perhaps, such ROS may even act on adjacent glia, potentially via redox-sensitive glial proteins such as transient receptor potential melastatin 2 (TRPM2), which have been shown to modulate synaptic plasticity (Wang et al., 2016; Turlova et al., 2018).

### 4.2 Downstream effector pathways of ROS and aquaporin-dependent structural plasticity

ROS can regulate the activity of several protein kinases, including those implicated in canonical neurodevelopmental pathways, either via modification of reactive amino acid residues on kinases or, indirectly, by redox-mediated inhibition of counteracting phosphatases (Finkel and Holbrook, 2000; Corcoran and Cotter, 2013; Holmström and Finkel, 2014). Notably, previous work has implicated aquaporin channels in modulating the efficacy of ROS-regulated protein kinase signalling. By altering a cell’s permeability to extracellularly-generated ROS, aquaporins can amplify or diminish the strength of redox-dependent downstream pathways (Miller et al., 2010; Bertolotti et al., 2013). For instance, mammalian Aquaporin 8, which shares ~33% of amino acid sequence identity with the *Drosophila* aquaporin channels, controls the entry of NADPH-oxidase derived H_2_O_2_ to increase growth factor signalling in human leukaemia B-cells (Vieceli Dalla Sega et al., 2017). Of particular interest are CaMKII and PKA signalling, given that their persistence can be enhanced be elevated cytosolic ROS (Humphries et al., 2007; Anderson, 2011), and that both pathways act to limit the elaboration of dendritic arbors in an activity-dependent manner. For example, targeted inhibition of CaMKII or PKA in otherwise non-manipulated neurons results in dendritic over-growth and an increase in arbor size and complexity (Wu and Cline, 1998; Zou and Cline, 1999; Tripodi et al., 2008). This is similar to what we have seen following quenching of extracellular H_2_O_2_ or knockdown of the aquaporins Bib and Drip, suggesting that either CaMKII or PKA signalling might be downstream effectors of activity regulated, Duox-generated extracellular H_2_O_2_.

Given the increasing number of signals involved in anterograde and retrograde signalling between neurons one might ask how ROS contribute to these signalling pathways? It is possible that Duox acts as an integrator of multiple signalling pathways, in that its activity is regulated by a number of pathways, including the Rho GTPase Rac1 (Hordijk Peter L., 2006) and calcium, via its EF-hands (Kawahara et al., 2007). It will be interesting to determine the range and temporal dynamics of Duox activity following neuronal activation; whether Duox reports on low, medium or high levels of neuronal activation, brief bursts or only following prolonged activation. Thus, it is conceivable that different inter-neuronal signalling pathways are utilised for distinct contexts, in terms of their activation pattern and, equally, their spatio-temporal dynamics of signalling

## Acknowledgements

The authors would like to thank members of the Landgraf lab for feedback on the manuscript. We are grateful to Andreas Bergmann, Paul Garrity, Won-Jae Lee, Paul Martin, Helen Weavers, Will Wood, as well as the Bloomington Drosophila Stock Center for generously providing fly stocks. This work was supported by funding from the Biotechnology and Biological Sciences Research Council to M.L. (BB/R016666/1). The work benefited from the Imaging Facility, Department of Zoology, supported by Matt Wayland and funds from a Wellcome Trust Equipment Grant (WT079204) and contributions by the Sir Isaac Newton Trust in Cambridge, including Research Grant (18.07ii(c)).

